# Effects of pharmacological modulators of α-synuclein and tau aggregation and internalization

**DOI:** 10.1101/2020.01.27.921643

**Authors:** Antonio Dominguez-Meijide, Eftychia Vasili, Annekatrin König, Maria-Sol Cima-Omori, Andrei Leonov, Sergey Ryazanov, Markus Zweckstetter, Christian Griesinger, Tiago F. Outeiro

## Abstract

Parkinson’s disease (PD) and Alzheimer’s disease (AD) are common neurodegenerative disorders of the elderly and, therefore, affect a growing number of patients worldwide. Both diseases share, as a common hallmark, the accumulation of characteristic protein aggregates, known as Lewy bodies (LB) in PD, and neurofibrillary tangles in AD. LBs are primarily composed of misfolded α-synuclein (aSyn), and neurofibrillary tangles are primarily composed of tau protein. Importantly, upon pathological evaluation, most AD and PD/Lewy body dementia cases exhibit mixed pathology, with the co-occurrence of both Lewy bodies and neurofibrillary tangles, among other protein inclusions. Recent studies suggest that both aSyn and tau pathology can spread and propagate through neuronal connections. Therefore, it is important to investigate the mechanisms underlying aggregation and propagation of these proteins for the development of novel therapeutic strategies. Here, we assessed the effects of different pharmacological interventions on the aggregation and internalization of tau and aSyn. We found that anle138b and epigallocatechin gallate decrease aSyn aggregation, that fulvic acid attenuates aggregation of the repeat domain of tau, and that dynasore reduces tau internalization. Establishing the effects of small molecules with different chemical properties on the aggregation and spreading of aSyn and tau will be important for the development of future therapeutic interventions.

## Introduction

Parkinson’s disease (PD) and Alzheimer’s disease (AD) are the most common neurodegenerative disorders and their prevalence is increasing due to the increase in aged population ^1^. Regardless of the differences in their clinical features, both are characterized by the accumulation of protein aggregates in the brain ^2–5^. One of the characteristics of these aggregates is the presence of misfolded forms of specific proteins. Thus, the presence of aggregates of misfolded α-synuclein (aSyn) is considered a hallmark of PD and aggregates of misfolded tau and amyloid-beta peptide in the brain are a hallmark of AD ^2, 5^. Recent studies suggest that aggregates of these proteins may spread throughout the brain propagating the disease ^6, 7^. Once the misfolded form of the protein enters into a new cell, it may act as a template for the misfolding of the natively folded form of the protein ^8–10^, leading to a loss of function and effectively propagating the disease ^11^. Therefore, the study of the molecular mechanisms underlying the aggregation and propagation of these proteins is important for the understanding of these diseases and the development of suitable therapeutic strategies.

Several different types of assays have been developed to study protein interactions in different organisms, amongst them, protein fragment complementation assays (PCAs) ^12^. In these assays, a fluorescent protein or an enzyme is truncated and fused to two proteins of interest. When these two proteins interact with each other, both complementary fragments lead to the reconstitution of the reporter activity, be it enzymatic or fluorescent. One of the PCA methods is bimolecular fluorescence complementation (BiFC), based on the reconstitution of a fluorescent protein upon the interaction of two proteins ^12–14^. The BiFC signal can be detected, for example, by fluorescence microscopy or by flow cytometry without the need for any other treatment to the cells. Using the BiFC principle, it is possible to monitor protein release and transfer upon co-culturing cells expressing the proteins of interest fused to each of the non-fluorescent fragments of the fluorescent protein.

In recent years, several different aggregation and internalization inhibitors have been identified, such as the aSyn aggregation inhibitors epigallocatechin gallate (EGCG) ^15–17^ and anle138b ^18^, and the tau aggregation inhibitor fulvic acid ^19, 20^. EGCG is a catechin present in green tea that redirects β-sheet rich amyloid into unstructured aSyn ^15, 16^. Nevertheles, clinical trials using EGCG were negative for reasons that are still not understood but, most probably, due to low bioavailability ^21^.

Anle138b is a diphenylpyrazole that reduces aSyn aggregation in various models of PD^18, 22^ and MSA ^23^, aggregation of amyloid beta (AB) in AB dependent mouse models of AD^24^ and aggregation of tau in tau-dependent mouse models of AD ^25, 26^, as well as aSyn spreading in a PD mouse model^18^.

Fulvic acid is a mixture of different polyphenolic acids produced by humus that attenuates heparin-induced tau aggregation *in vitro* ^19^. On the other hand, the dynamin inhibitor dynasore inhibits clathrin-dependent endocytosis. Dynamin is a protein that cleaves the fusion between the cell wall and the vesicle formed in clathrin-mediated endocytosis, which is required for the internalization of the vesicle ^27^.

The versatility of the BiFC system enables the assessment of the effects of genes and small molecules on the aggregation of proteins such as aSyn and tau. Here, we assessed the effects of EGCG, anle138b, fulvic acid and dynasore on the aggregation and internalization of aSyn and tau *in vitro* and using the BiFC assay using human cells as a living test tube.

## Results

### aSyn and tau are released to cell culture media and taken up by neighboring cells

First, we assessed the release and uptake of aSyn and tau in different cell types. We used the BiFC system with Venus fluorescent protein as the reporter fluorophore. Co-culturing cells expressing the protein of interest (aSyn or tau) fused to either VN- or VC-fragments enabled us to assess, simultaneously, the release and uptake of the proteins, as it requires that at least one of the constructs is released and taken up by neighboring cells. Once the proteins are released and taken up, the fluorophore is reconstituted in the recipient cells and fluorescence is emitted (Fig. 1A). Using flow cytometry, we found 1 ± 0.5 % of BiFC positive cells for aSyn and 3.9 ± 1.4 % of BiFC positive cells for tau. These values, though relatively small, were different from those obtained for the negative controls (Fig. 1). When both constructs were co-expressed in the same cell, the fluorescence values were similar to those obtained with GFP (Supplementary Figure 1).

**Figure 1.**
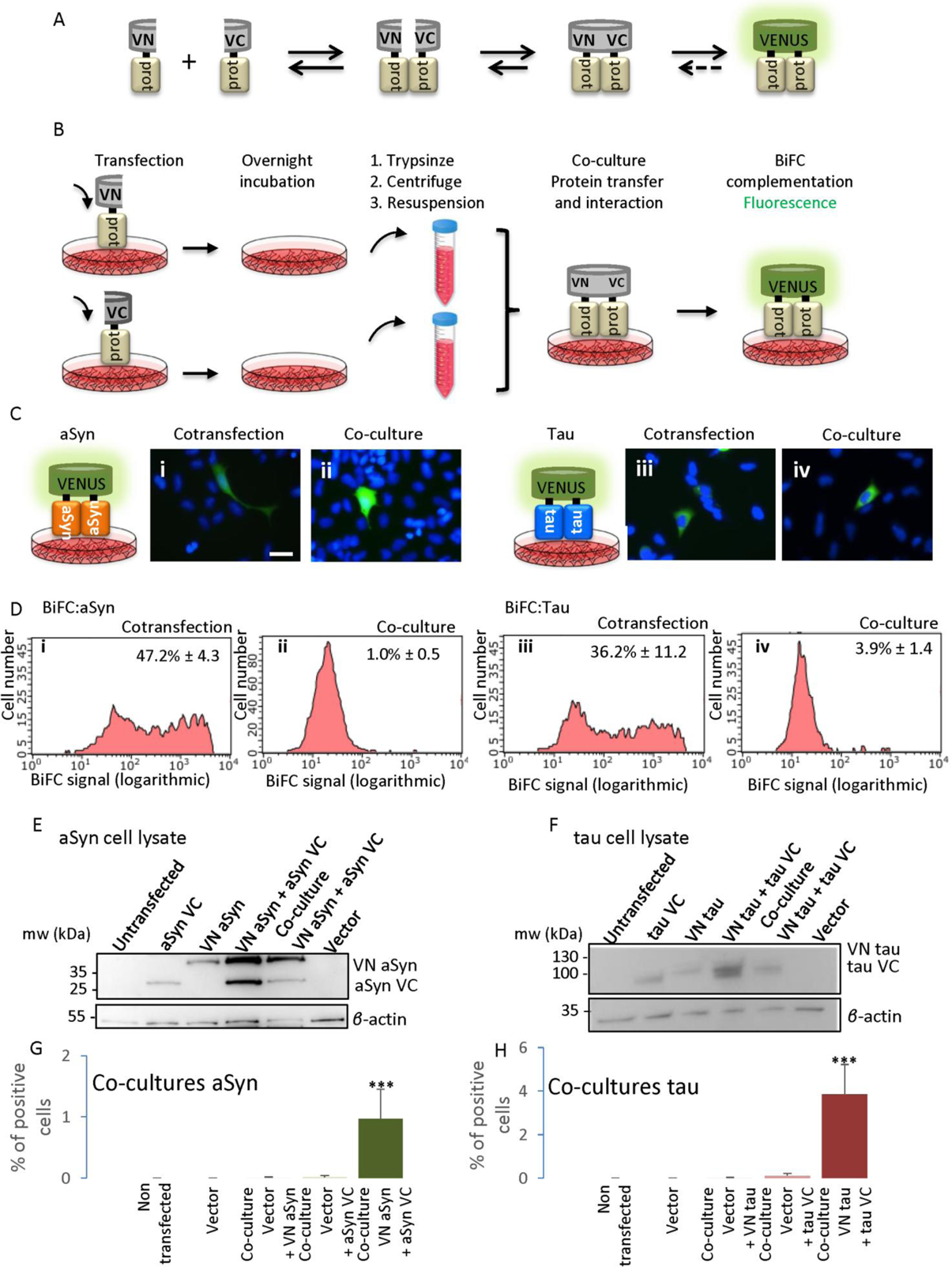
Reconstitution of venus fluorescence upon co-culture of transfected cells indicates release and uptake of aSyn or tau proteins. A. Representative scheme of the BiFC principle. Proteins tagged to the VN fragment of the venus protein interact with proteins tagged to the VC fragment of the venus protein leading to the reconstitution of the fluorophore. B. Schematic of the co-culture process. Cells transfected with proteins tagged with the VC fragment of venus were trypsinized and mixed with cells transfected with proteins tagged to the VN fragment. Protein release and uptake by new cells results in an interaction and reconstitution of the fluorophore. C. Epifluorescence microscopy showing positive cells. i) Representative picture of cells cotransfected with aSyn BiFC constructs. ii) Representative picture of aSyn co-cultured cells. iii) Representative picture of cells cotransfected with tau BiFC constructs. iv) Representative picture of tau co-cultured cells. Scale bar 50 µm. D. Flow cytometry analysis of different cultures. i) Representative histogram showing the number of cells versus fluorescence intensity for aSyn cotransfected cells. ii) Representative histogram showing the number of cells versus fluorescence intensity for aSyn co-cultured cells. iii) Representative histogram showing the number of cells versus fluorescence intensity for tau cotransfected cells. iv) Representative histogram showing the number of cells versus fluorescence intensity for tau co-cultured cells. The average percentage of positive cells ± SD is shown in each histogram. Cells with a value of fluorescence intensity above 120 fluorescence units are considered positive. E. Western blot showing the levels of all aSyn constructs. Full length blots are presented in Supplementary Figure 6A. F. Western blot showing the levels of all tau constructs. Full length blots are presented in Supplementary Figure 6B. G. Percentages of positive cells in co-cultures are significantly different from those obtained for the negative controls for the aSyn constructs. H. Percentage of positive cells in co-cultures are significantly different from the percentage of positive cells in the negative controls for the tau constructs. *** p < 0.0005 from all other groups. ANOVA and subsequent paired t-test. Error bars represent SD.

To confirm that all expressed proteins were released, we performed western blot analyses (Fig. 2A-B; Supplementary Figure 2) and ELISA of the conditioned media in which cells were growing (Fig. 2C-D).

**Figure 2.**
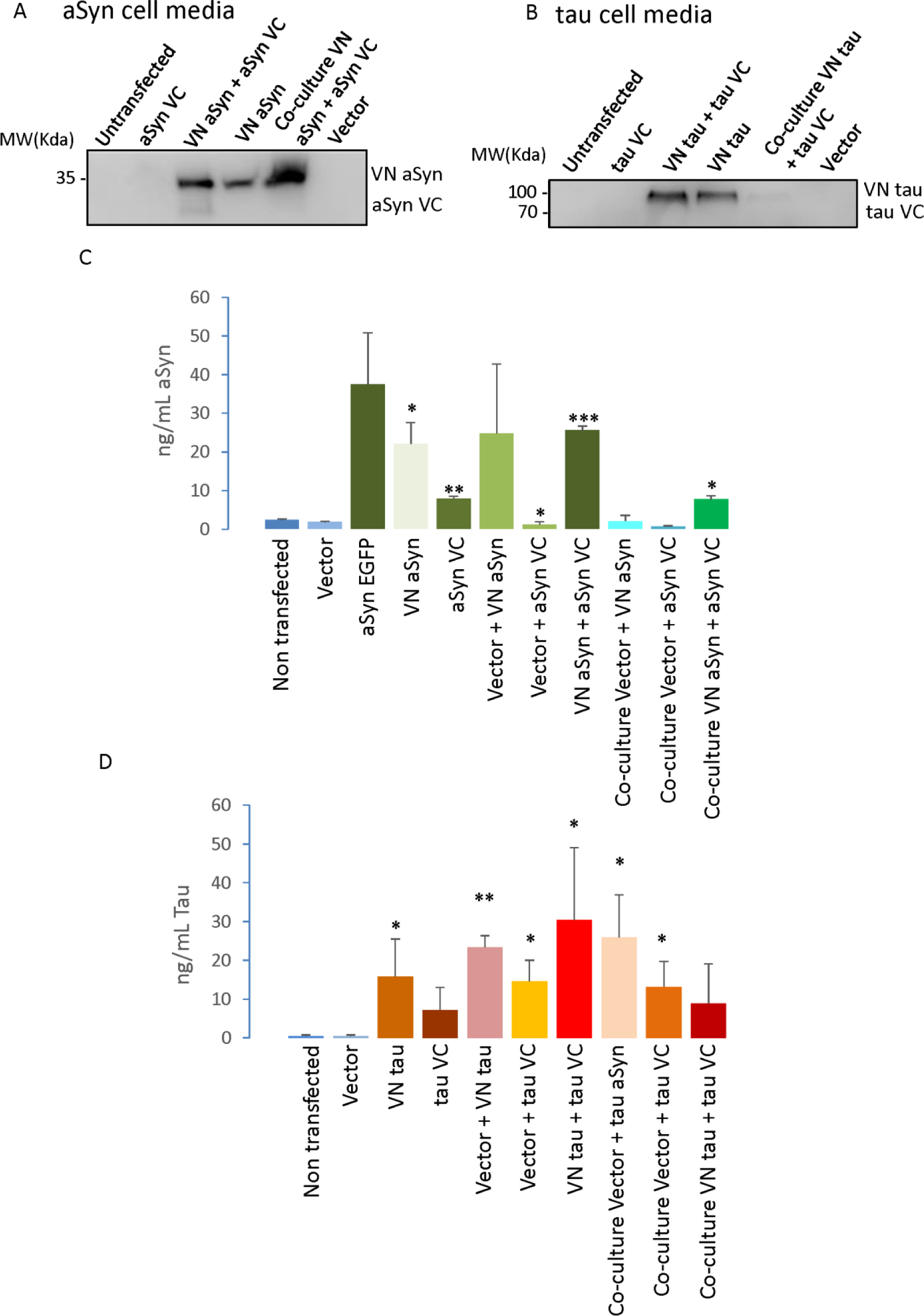
Proteins are released to the cell culture media. A. Western blot showing presence of different aSyn BiFC fragments in cell culture media. Full length blots are presented in Supplementary Figure 6C. B. Western blot showing presence of different tau BiFC fragments in cell culture media. Full length blots are presented in Supplementary Figure 6D C. ELISA measurements of aSyn levels in cell culture media show presence of the protein in cell media (N=4). D. ELISA measurements of tau levels confirm the presence of the protein in the cell culture media (N=5). * P < 0.05 ** p < 0.005 *** p < 0.0005 in relation to empty vector. Error bars represent SD. ANOVA and subsequent paired t-test in relation to empty vector.

Next, we added conditioned media from cells expressing one of the BiFC constructs to cells expressing the complementary BiFC construct, in order to determine whether the different proteins were released and taken up (Fig. 3A). We confirmed that addition of conditioned media induced the reconstitution of Venus fluorescence in all cases, confirming that both aSyn and tau can be released and taken up by cells (Fig. 3B). This prompted us to subsequently investigate the effect of the internalization inhibitor dynasore. Additionally, in the case of tau, both sarkosyl-solube and sarkosyl-insoluble fractions are detected in the media, suggesting that not only monomeric, but also aggregated forms of tau may be released from cells (Supplementary Figure 2) ^28, 29^.

**Figure 3.**
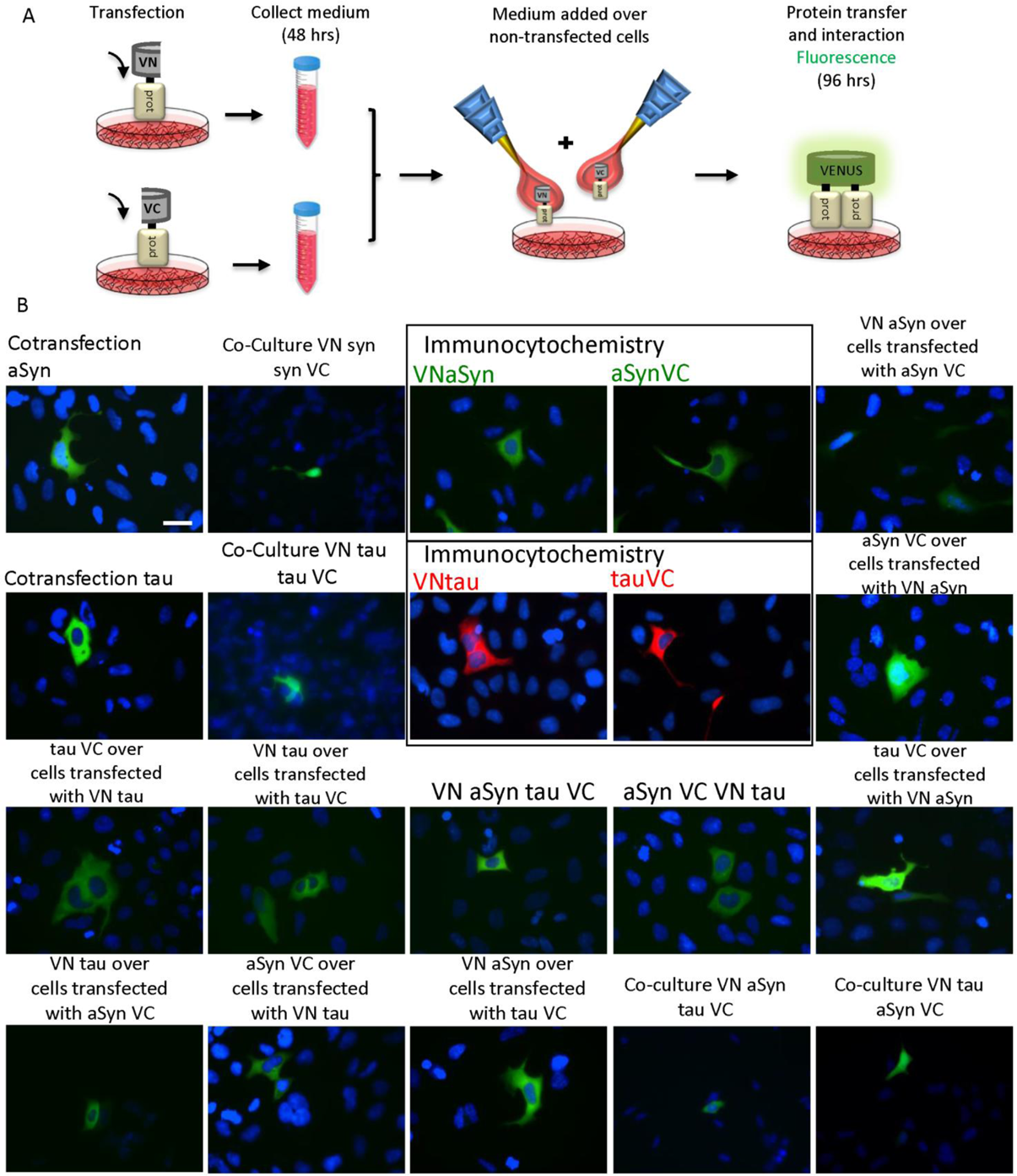
Conditioned media added to cells reports on protein release and uptake. A. Schematics of the experimental setup. Cells are transfected with the different constructs, after 48 hours, media are collected and added to either non-transfected cells or to cells transfected with a different construct. The two proteins interact inside the receptor cells leading to reconstitution of the fluorophore and fluorescence. B. Representative pictures of the different combinations performed. In the negative cases, an immunocytochemistry was performed to assess the internalization of the proteins. Colors used for immunocytochemistry (pictures inside squares) are: green for aSyn and red for tau. Positive BiFC combinations yield green color. Mixes of aSyn and tau also yield geen color additionally showing the interaction amongst proteins. Scale bar 50 µm.

### Tau internalization is inhibited by dynasore

After assessing release and internalization, and based on the results above, we tested the effects of the dynamin inhibitor dynasore. Here, we focused only on tau internalization since aSyn internalization has already been studied through similar assays ^30, 31^. To assess the effect of dynasore on tau internalization, we initially confirmed the expression of dynamin 1 and dynamin 2 in HEK293 cells using immunocytochemistry (Fig. 4B-C). As previously reported, the expression of dynamin 1 is almost negligible in comparison with that of dynamin 2 ^32^. Next, we tested whether the dynamin inhibitor dynasore inhibited tau internalization in living cells, using the co-culture model described above (Fig. 1 and Fig. 4A). We observed a significant reduction (P = 0,027) in the percentage of fluorescent cells upon treatment with dynasore (Fig. 4E), but no effect on fluorescence intensity or in protein levels (Fig. 4F-H). These findings, in combination with the results mentioned above, suggest that the reduction in the percentage of fluorescent cells was due to the inhibition of protein internalization.

**Figure 4.**
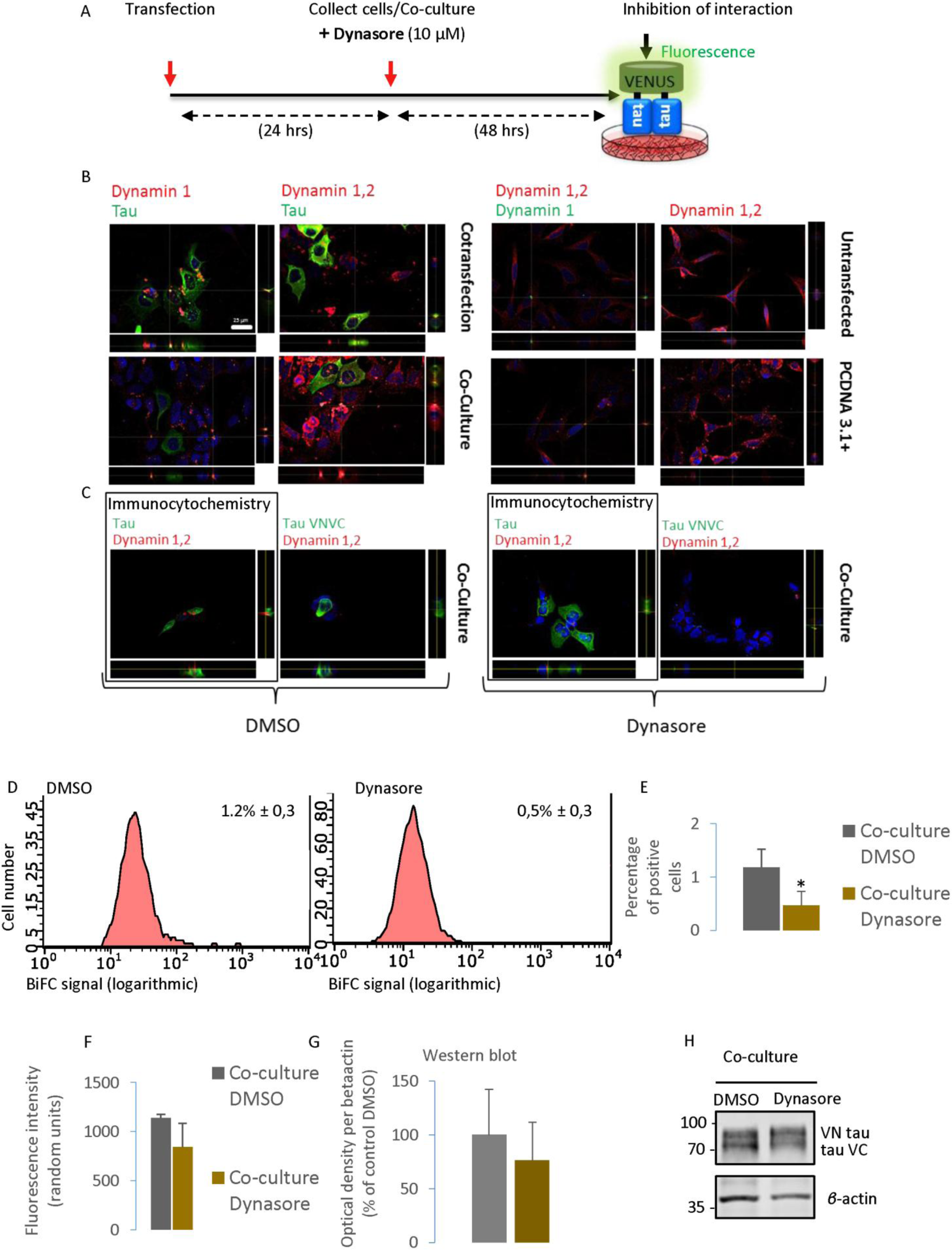
Effect of inhibition of tau internalization. A. Schematic representation of the experimental setup. Cells were co-cultured and treated with 10 µM dynasore. Inhibiting this internalization should lead to no fluorescence. B. Representative immunofluorescence pictures showing the expression of dynamin 1 and dynamin 2. Both dynamin 1 and 2 are detected in HEK293 cells by immunocytochemistry. Scale bar 25 µm. C. Representative pictures of vehicle and dynasore-treated co-cultures. Tau is present in dynasore-treated cells. D. Flow cytometry results for vehicle and dynasore-treated co-cultures showing the average percentages of positive cells ± SD. E. Flow cytometry results show that dynasore-treated co-cultured cells show a significant decrease in percentage of positive cells in comparison with vehicle-treated co-cultured cells (N=4). F. Flow cytometry results show that dynasore-treated co-cultured cells show no significant decrease in fluorescence intensity in comparison with vehicle-treated co-cultured cells (N=4). G. Western blot results show that dynasore-treated co-cultured cells show no significant decrease in protein expression levels in comparison with vehicle-treated co-cultured cells (N=3). H. Western blot picture of the vehicle-treated and dynasore-treated co-cultured cells. Full length blots are presented in Supplementary Figure 6 E. Data are shown as mean ± SD. * p < 0.05 t-test.

### EGCG and anle138b reduce aSyn aggregation *in vitro*

Initially, we assessed the effect of EGCG and anle138b on aSyn (10 μg/ml) aggregation *in vitro*, using RT-QuiC ^33–35^. We found that treatment with either EGCG (10 nM) or anle138b (100 nM) reduced ThT fluorescence intensity, confirming that both compounds reduced aSyn aggregation (Fig. 5G, K). In addition, we found that both substances lead to a significant decrease (P = 0,012 for anle138b and P = 0,009 for EGCG) in the amount of monomers incorporated per amplification cycle ^36, 37^ when compared to the control (Fig. 5H, L).

**Figure 5.**
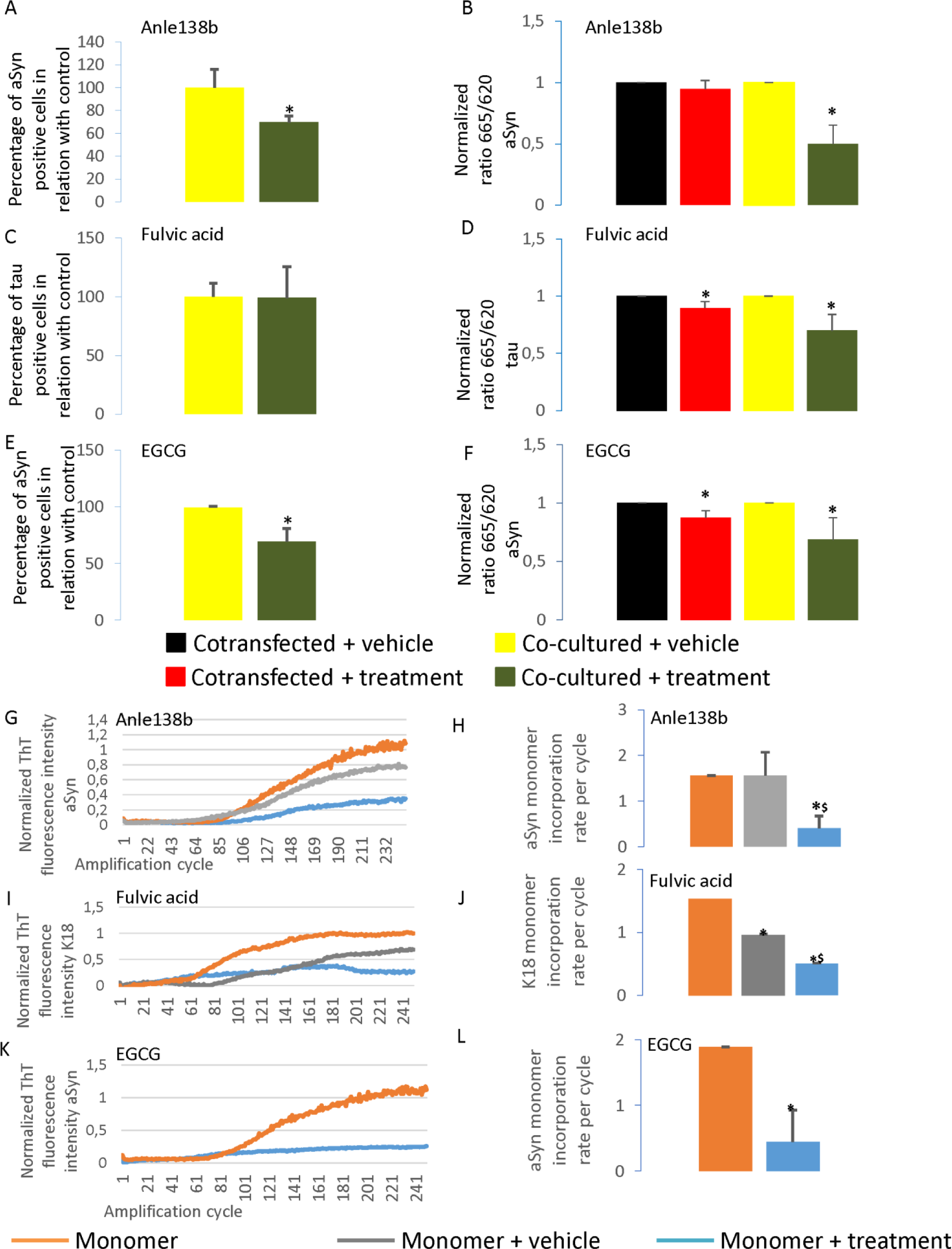
Anle138b, fulvic acid and EGCG inhibit protein aggregation. A. Flow cytometry results of co-cultured cells treated with anle138b (1 µM) show a significant decrease in the percentage of positive cells in comparison with cells treated with vehicle (N = 3). B. HTRF measurements of aSyn-transfected and co-cultured cells treated with the aggregation inhibitor anle138b (1 µM) show significant differences in aggregation (N = 4). C. Flow cytometry results of cells treated with the aggregation inhibitor fulvic acid (37 μM) show no significant differences in the percentage of positive cells in relation to cells treated with vehicle (N = 3). D. HTRF measurements of Tau-transfected cells treated with the aggregation inhibitor fulvic acid (37 μM) show significant differences in aggregation (N = 4) E. Flow cytometry results of co-transfected cells treated with the aggregation inhibitor EGCG (0,1 µM) show statistically significant differences in the percentage of positive cells in relation to cells treated with vehicle (N = 3). F HTRF results of aSyn-transfected cells treated with the aggregation inhibitor EGCG (0,1 μM) show significant differences in aggregation (N = 4). G. RT-QuiC graph of the monomeric aSyn amplification in the presence and absence of 100 nM anle138b (N=4). Protein amplified in presence of 100 nM anle138b shows lower ThT fluorescence intensity than in absence of anle138b. Orange: only monomeric protein added; grey: monomeric protein and DMSO added; blue: monomeric protein and anle138b (dissolved in DMSO) added. H. RT-QuiC results show a significant decrease in monomer incorporation rate per cycle when amplification in the presence of 100 nM anle138b. Orange: only monomeric protein added; grey: monomeric protein and DMSO added; blue: monomeric protein and anle138b (dissolved in DMSO) added. I. RT-QuiC graph of the tau MBD (K18) amplification in the presence and absence of 37 µM fulvic acid (N=4). Protein amplified in presence of fulvic acido shows lower ThT fluorescence intensity than in the absence of fulvic acid. Orange: only monomeric protein added; grey: monomeric protein and methanol added; blue: monomeric protein and fulvic acid (dissolved in methanol) added. J. RT-QuiC assay shows a significant decrease in monomer incorporation rate per cycle in the presence of 37 µM fulvic acid. Orange: only monomeric protein added; grey: monomeric protein and DMSO added; blue: monomeric protein and anle138b (dissolved in DMSO) added. K. RT-QuiC graph of the monomeric aSyn amplification in the presence and absence of 10 nM EGCG. Protein amplified in presence of 10 nM EGCG shows lower ThT fluorescence intensity than in absence of EGCG. Orange: only monomeric protein added; blue: monomeric protein and EGCG added. L. RT-QuiC results show a significant decrease in monomer incorporation rate per cycle in the presence of 10 nM EGCG (N=4). Orange: only monomeric protein added; blue: monomeric protein and EGCG added. * p< 0.05 in comparison with control-treated cells. Error bars represent SD.

### Fulvic acid reduces tau K18 aggregation *in vitro*

Next, we assessed the effect of fulvic acid (37 µM) on tau (7 µg/ml) aggregation *in vitro*, using RT-QuiC. Since the fragment of tau that is mostly responsible for its aggregation is the microtubule binding domain (MBD), we performed the RT-QuiC amplification using the MBD of 4-repeat tau, termed K18 ^38^. We found that fulvic acid reduced the ThT fluorescence intensity, suggesting reduced aggregation of tau K18. Additionally, we observed a decrease (P = 0,0023) in the number of monomers incorporated per amplification cycle (Fig. 5I-J) when we used fulvic acid at its IC50 ^19^.

### Anle138b reduces aSyn aggregation in living cells

Next, we assessed whether anle138b also affected aSyn aggregation the context of a human cell line (HEK293). We transfected cells with the BiFC constructs and performed co-cultures as described. 24 hours after co-culturing, cells were treated with anle138b or with vehicle only, as a control. 12 hours after treatment cells were collected and fluorescence was assessed by flow cytometry. Absence of interaction should lead to absence of fluorescence, and lower fluorescence percentages. We observed no changes in fluorescence intensity nor in the total percentage of fluorescent cells on the whole population analyzed (Supplementary Figure 3F and H). As this could be due to high variability between samples, we normalized the results and expressed them as relative values using 100% for cells treated with vehicle (relative percentage of positive cells in relation to the control). Interestingly, when we compared the relative percentage of positive cells in relation to the control, we found a significant decrease (decrease to 69,71 ± 5,27%, P = 0,037), suggesting inhibition of aggregation (Fig. 5 A-B and Supplementary Figure 3). To further confirm this, we assessed aggregation using a commercially-available HTRF-based assay that measures protein aggregation, based on the Förster resonance energy transfer principle. The ratio between the fluorescence emission measured at 665 nm (FRET) and the emission measured at 620 nm (emission of FRET donor) indicates a high proximity between proteins. As such, higher 665/620 ratios indicate higher aggregation. Due to the high variability, we normalized the results obtained for each sample to its corresponding control. When comparing co-cultured cells treated with anle138b with co-cultured cells treated with vehicle, we found a significant decrease (P = 0,007) in the normalized 665/620 ratio ^39, 40^ (a decrease to 49,91 ± 15,30% (Fig. 5B, green bar)), confirming the reduction in aggregation. Interestingly, we found no significant differences in cells transfected with both constructs, suggesting that anle138b did not disaggregate aSyn aggregates but, instead, inhibited the aggregation process (Fig. 5B, red bar). Finally, we confirmed that anle138b did not affect the levels of aSyn (Supplementary Figure 3).

### EGCG decreases aSyn aggregation in living cells

Next, we performed and identical study with EGCG, an established aggregation inhibitor. The co-cultures were performed as described above and treated with EGCG for 12 hours. Then, cells were collected and processed for flow cytometry. We observed no changes in fluorescence intensity nor in total percentage of fluorescent cells (Supplementary Figure 5F and H). Interestingly, when we compared the relative percentage of positive cells in relation to the control, we observed a significant decrease (P = 0,0107) upon EGCG treatment (a decrease to 69,31 ± 11,44%), suggesting an inhibition of the aggregation process (Fig. 5E and Supplementary Figure 5).

To further confirm this result, we used the HTRF-based aggregation assay. Again, we found a decrease in the normalized 665/620 fluorescence ratio in cells treated with EGCG (decreases to 87,43 ± 5,84, P = 0,023 (Fig. 5F, red bar) and 68,51± 18,83%, P = 0,044 (Fig. 5F, green bar), respectively), suggesting that this compound may disaggregate aSyn aggregates and inhibit the aggregation process (Fig. 5F). Importantly, we also found that EGCG did not affect the protein levels, supporting the direct effect on aggregation (Supplementary Figure 5).

### Fulvic acid reduces tau aggregation in living cells

Next, a similar approach was used to study the effect of fulvic acid on tau aggregation. We transfected cells with tau BiFC constructs and performed the co-cultures as described. 24 hours after co-culturing, cells were treated with fulvic acid and 12 hours after treatment cells were collected and processed for flow cytometry. We observed no changes in fluorescence intensity, in total percentage of fluorescent cells or in the relative percentage of positive cells in relation with control (decrease to 99,05 ± 26,50%) (Fig. 5C and Supplementary Figure 4).

Interestingly, when we assessed aggregation using a tau-specific HTRF-based assay, we observed a significant decrease in the normalized 665/620 ratio, both in cells expressing both constructs and in co-cultures (decrease to 89,73 ± 5,43%, P = 0, 032 (Fig. 5D, red bar) and 74,14, P = 0,041 ± 15,01% (Fig. 5D, green bar), respectively) (Fig. 5D and Supplementary Figure 4). Again, we also found no effect of fulvic acid on the levels of tau (Supplementary Figure 4).

## Discussion

Neurodegenerative disorders, such as PD and AD, are characterized by misfolding and aggregation of proteins in specific brain regions. Whether this protein aggregation is causative or an epiphenomenon is still unclear, although growing evidence suggests protein aggregation causes cellular and circuit dysfunction that may, ultimately, result in cell death. Therefore, the study of the molecular mechanisms underlying protein aggregation and transference of these proteins between cells is of utmost importance for the development of novel therapeutic strategies. In this context, modulation of protein aggregation has been an attractive hypothesis. In PD, inhibition of aSyn aggregation is seen as a possible strategy ^41^. Likewise, the initial results of the aducanumab trial in AD showed no cognitive benefit, but more recent analyses showed a possible benefit at the highest dose ^42, 43^. This shows the need for further understanding the underlying mechanisms of toxicity, and the need to develop alternative therapeutic approaches ^42^. In this regard, inhibition of tau aggregation may be a possible therapeutic option ^25, 26, 44, 45^. Here, we assessed the effects of aggregation modulators for direct comparison: anle138b and EGCG for aSyn, and fulvic acid for tau.

Using simple, yet powerful, human cell-based models, we found that anle138b reduces aSyn aggregation in a HEK293 venus BiFC cell model. Until now, anle138b was mostly tested in mouse models ^18, 23, 46^. In cells, it was tested in melanoma and in H4 cells ^47, 48^. These previous studies, together with our results, show the ability of the compound to decrease aSyn aggregation, both in cells and in vitro, as assessed by RT-QuiC. Interestingly, the lack of effect observed in a BiFC-based aSyn assay, suggests that, in this model, anle138b may not be able to dissolve previously formed oligomers/aggregates. This is consistent with studies in which treatment with anle138b was performed before RT-QuiC amplification ^49^, and with the proposed mechanism of action of the compound on aSyn aggregation, which is thought to be based on the direct targeting of oligomeric species, blocking interpeptide interactions and preventing the spontaneous formation of ordered beta-sheet structures ^18, 50^. On the other hand, it has been reported that, in mouse brain, anle138b reduces the density of aSyn aggregates while increasing the number of monomeric and small assemblies of truncated aSyn ^46^. Nonetheless, in transgenic aSyn mouse model using the PLP promoter, once pathology is too advanced and mainly induced by a neurotoxin, there were no significant effects of anle138b on neuronal loss or in the density of intracytoplasmatic aSyn inclusions, in agreement with our results ^51^. In the MI2 model, results were observed at a stage where motor function was already impaired, showing that, in some animal models, anle138b has a curative effect. Such curative effects were also observed for Ab ^24^ and tau based ^25^ AD models. It is possible that the differences in the effects observed are due to intrinsic differences in the models – HEK cells vs. neurons in the context of the organ in an animal and the duration of the treatment (several weeks in animals, few hours in cells). Nevertheless, both types of models agree that the effects are not simply a result from changes in the levels of aSyn ^18^. According to this proposed mechanism of action, anle138b also appears effective in other proteinopathies ^18, 24–26, 49, 52^, and it has been shown to alleviate motor deficits in a mouse model of MSA ^23, 51^.

Our work also validates the effect of EGCG, traditionally obtained from green tea leaves, as an aSyn aggregation inhibitor ^16, 17^. This has been observed *in vitro*, using ThT-based aggregation assays ^17, 53, 54^, and in cells, in OL-93 and AS-PC12 cells ^53, 55^. Our findings are consistent with the previous reports, and extend the previous observations to different cell models and using different readouts of aggregation. In particular, our data suggest two possible activities: in preventing aggregation and also in disaggregating previously formed inclusions. This is consistent with previous reports in which treatment with EGCG was performed in already formed aggregates ^16, 17, 56^. This effect was not only reported for aSyn, but also for other amyloidogenic proteins, such as AB and γ-synuclein (gSyn) ^16, 56, 57^. Such effects are in agreement with the proposed mechanism of action for EGCG, whereby it mediates the reorientation of bonds between ordered protein molecules, leading to amyloid remodeling and the appearance of unordered, amorphous protein aggregates ^17, 58^. We also investigated the effect of fulvic acid on tau aggregation, using our cell models. Fulvic acid is a mixture of polyphenolic acid compounds produced by humus ^59^, and was shown to reduce the aggregation of amyloid β or the prion protein ^60, 61^. Our results showed that the compound inhibits K18 tau aggregation *in vitro*, and of full length tau in a cell model. Interestingly, fulvic acid seems to disaggregate previously formed tau aggregates in cells, and is consistent with findings using heparin-induced tau aggregation ^19^.

Finally, we assessed whether blocking endocytosis would alter aggregation, possibly by reducing the seeding step of protein aggregation ^8, 62^. Dynamins play essential roles in membrane remodeling and fission of clathrin-coated vesicles formed during endocytosis ^63, 64^. Dynamin polymerizes to form a helix around the neck of budding vesicles of plasma membrane leading to membrane fission and generation of free clathrin-coated vesicles ^64, 65^. Dynasore is a non-competitive, reversible dynamin1 and dynamin2 inhibitor that interferes with the catalytic step of the GTPase activity of the proteins, leaving GTP bound to dynamin2 and dynamin1 ^65, 66^. Treatment with dynasore has two different effects: (i) the blocking of the detachment of fully-formed coated pits from the membrane; and (ii) the reduction in plasma membrane cholesterol, thereby reducing the fluidity of the cell membrane ^64, 65, 67, 68^. Dynasore interferes with aSyn uptake in primary human adult neurons ^69^, but it has been shown in H4 cells that this does not happen through clathrin mediated mechanisms ^31^. Our study shows that, dynasore reduces the percentage of BiFC-positive cells in co-cultured cells, without changing the fluorescence intensity or protein levels, consistent with a reduction in tau uptake.

In summary, we found that anle138b and EGCG reduce aSyn aggregation. We also found that fulvic acid reduces tau aggregation, and that dynasore blocks tau endocytosis, providing further validation for the effects of these molecules and their targets as relevant agents in the context of synucleinopathies and tauopathies.

## Methods

### Recombinant protein preparation

Recombinant aSyn was prepared as described before ^70^. In brief, pET21-asyn was transformed into competent BL21-DE3 cells (Sigma). Bacteria were then incubated at 37°C in 2x LB medium to an OD_600_ of 0.6. Protein expression was induced with 1mM isopropyl-β-D-1-thiogalactopyranoside (Peqlab) for 3 hours. Cells were harvested by centrifugation (6000g, 20 min). After centrifugation, cells were lysed (750 mM NaCl, 10 mM Tris, 1 mM EDTA pH 7.6, protease inhibitor (Roche, Mannheim, Germany)), sonicated and incubated at 95°C for 15 min. Cell lysates were then centrifuged (15000 g, 20 min) ant the supernatants were collected and dialysed (1mM EDTA, 50 mM NaCl, 10 mM TRIS pH 7.6). After dyalisis, supernatants purified with anion exchange chromatography (HiTrap Q HP, GE Healthcare) using a mobile phase of 25 mM Tris pH 7.7, followed by size exclusion chromatography (HiLoad Superdex 75, GE Healthcare). Following purification, protein was concentrated using a centrifugation filter (3K, Amicon, Merck, Darmstadt, Germany) up to a final concentration of 5 mg/mL and stored at −80°C.

K18 tau was prepared as described previously ^71^. In brief, human tau constructs were expressed in the vector pNG2 (Merck-Novagen, Darmstadt) in *E. coli* strain BL21(DE3). Expressed proteins were then purified from bacterial extracts by making use of the protein heat stability and subsequent FPLC SP-Sepharose chromatography (Amersham Biosciences). The cell pellets were resuspended in boiling extraction buffer (50 mm MES, 500 mm NaCl, 1 mm MgCl2, 1 mm EGTA, 5 mm dithiothreitol, pH 6.8) complemented with a protease inhibitor mixture. Following this, cells were disrupted with a French pressure cell and subsequently boiled for 20 min. The soluble extract was isolated by centrifugation, the supernatant was dialyzed against two changes of cation exchange chromatography buffer A (20 mm MES, 50 mm NaCl, 1 mm EGTA, 1 mm MgCl2, 2 mm dithiothreitol, 0.1 mm phenylmethylsulfonyl fluoride, pH 6.8) and loaded into an FPLC SP-Sepharose column. The proteins were eluted by a linear gradient of cation exchange chromatography buffer B (20 mm MES, 1 m NaCl, 1 mm EGTA, 1 mm MgCl2, 2 mm dithiothreitol, 0.1 mm phenylmethylsulfonyl fluoride, pH 6.8). NMR samples contained 0.9–1.5 mm 15N- or 15N/13C-labeled protein in 95% H2O, 5% D2O, 50 mm phosphate buffer, pH 6.8, with 1 mm dithiothreitol.

### HEK293 cell line culture

Human embryonic Kidney (HEK293) cells were cultured in DMEM with 10% FBS and 1% penicillin/streptomycin at 37°C and 5% CO_2_ in a humidified incubator. Twenty-four hours prior to transfection approximately 100.000 HEK293 cells were plated per well in a 12-well plate (Costar, Corning, New York, USA). Transfection was performed with Metafectene® according to the following protocol: 1,5 µg of total DNA were added to 50 µl of DMEM medium without adds and this mixture was added to a solution containing 3µl of Metafectene in 50 µl of DMEM. The resulting mixture was added dropwise to the cells and the plate was gently rocked.

Sixteen hours after transfection HEK293 cells were fed with fresh medium and the co-cultures were performed as follows: HEK293 cells transfected with the empty vector (PCDNA3.1+), aSyn VC, VN aSyn, Tau VC or VN Tau were trypsinized and cultured at a total density of 100.000 cells (50.000 coming from each transfection) per milliliter in different combinations.

Cells were kept at 37°C and 5% CO_2_ for an additional 48 hours.

### Anle138b treatment

Sixteen hours after transfection HEK293 cells were fed with fresh medium and co-cultured as described above. To ensure the transfer of proteins from one cell to another, cells were grown for 24 hours after co-culture and the presence of fluorescent cells was checked by microscopy before proceeding with further treatments. The following day, media was changed, new media without FBS was added and cells were treated with anle138b at a concentration of 1 µM ^18^. Twelve hours after treatment, cells were collected for flow cytometry, western blot and homogeneous time-resolved fluorescence (HTRF) ^39, 40^.

### EGCG treatment

Sixteen hours after transfection HEK293 cells were fed with fresh medium and co-cultured as described above. To ensure the transfer of proteins from one cell to another, cells were grown for 24 hours after co-culture and the presence of fluorescent cells was checked by microscopy before proceeding with further treatments. The next day, media was changed, new media without FBS was added and cells were treated with EGCG (Sigma-Aldrich, St. Louis, MO, USA) at a concentration of 0,1 µM. Twelve hours after treatment, cells were collected for flow cytometry, western blot and HTRF.

### Fulvic acid treatment

Sixteen hours after transfection HEK293 cells were fed with fresh medium and co-cultured as described above. To ensure the transfer of proteins from one cell to another, cells were incubated for 24 hours after co-culture and the presence of fluorescent cells was assessed by microscopy before proceeding with further treatments. The next day, media was changed, new media without FBS was added and cells were treated with fulvic acid (Cayman Chemical, Ann Arbor, MI, USA) at a concentration of 37 µM ^19^. Twelve hours after treatment, cells were collected for flow cytometry, western blot and HTRF.

### Dynasore treatment

Sixteen hours after transfection HEK293 cells were fed with fresh medium, co-cultured as described above and treated with dynasore (Sigma-Aldrich, St. Louis, MO, USA) to a final concentration of 10 µM. Twenty four and 48 hours after treatment, media was changed and the cells were treated with additional dynasore at 10 µM. Thirty minutes after the last treatment cells were collected for flow cytometry and western blot. Additionally some cell cultures were cultured on coverslips for immunocytochemistry.

### Flow cytometry

Cells were trypsinized, and trypsin was added to each plate and neutralized with media. The cell suspension was centrifuged, the supernatant discarded and the pellet was resuspended in DPBS with a 0,1 % of propidium iodide. 5000 events were counted in triplicates for each experiment in a Millipore Guava EasyCite flow cytometer.

### Sarkosyl extraction

Sarkosyl extraction was performed as described before ^72^. In brief, cells were trypsinized and washed with PBS to remove trypsin from the cell pellet. 500 µl of cell media was collected and the presence of adenylate kinase was measured with the ToxiLightTM cytotoxicity assay kit (Lonza, Basel, Switzerland) to check the integrity of the cell membrane. 750 µl of media were then collected and centrifuged for 5 minutes at 2000 rpm to get rid of cell debris. Cells were then resuspended in buffer (10 mM Tris HCl, 150 mM NaCl, 1 mM EGTA, 5 mM EDTA, 1% Sarkosyl, protease inhibitor as specified above, pH 7,4) while media was diluted 1:1 in 2x buffer. Cell lysates and media solutions were then incubated on ice for 15 minutes. For cell lysis, the suspension was syringe sheared 10 times with a 27G needle followed by incubation on ice for additional 15 minutes. Cells were there sonicated 2 times at 30% power and were incubated at 25°C for 20 minutes. For media preparation, media was centrifuged at 2000g and 4°C for 10 minutes, supernatants were then collected, and incubated on ice for an additional 15 minutes. All samples were then ultracentrifuged at 186000 g and 4°C for 60 minutes. Supernatants were saved as sarkosyl-soluble fraction and the pellet was resuspended in 100 µl of urea buffer (8 M Urea, 2% SDS, 50 mM Tris-HCl).

### Immunoblotting analyses of media and cell lysates

Cultured cells were lysed in RIPA buffer containing protease inhibitor cocktail (1 tablet/15 mL) (Roche Diagnostics, Mannheim, Germany), sonicated and stored at −80°C until analysis. Cell media was collected and the presence of adenylate kinase was measured with the ToxiLight^TM^ cytotoxicity assay kit (Lonza, Basel, Switzerland) to check the integrity of the cell membrane. 750 µl of media were then collected and centrifuged for 5 minutes at 2000 rpm to get rid of cell debris. Supernatants were collected and concentrated ten times in an Amicon 10 k centrifuge filter (Millipore Merck, Darmstadt, Germany) following the manufacturer’s instructions.

Protein concentrations in the lysates were determined by the Bradford protein assay and 12 µg of protein were loaded into 12% Bis-Tris polyacrylamide gel and transferred to nitrocellulose membranes. The membranes were blocked with 5% skim milk in TBS-Tween and incubated overnight at 4°C with primary antibodies, 1:3000 aSyn 1 antibody (BD Biosciences, San Jose, CA, USA), 1:6000 anti-total tau (DAKO Agilent, Santa Clara, CA, USA) and 1:10000 anti-b-actin (Sigma-Aldrich, St. Louis, MO, USA). After three washes with TBS-Tween, membranes were incubated for two hours with horseradish peroxidase (HRP) conjugated secondary antibodies (GE Healthcare, Helsinki, Finland) at 1:6000.

After incubation with the secondary antibody, membranes were washed three times with TBS-Tween and developed in a chemiluminescence system (Fusion FX Vilber Loumat).

### Immunoblotting of sarkosyl extracts

Protein concentration in sarkosyl extracted samples was determined using the 2D-Quant Kit (GE Healthcare) following the manufacturer’s instructions and 12 µg of protein were loaded into 4-15% gradient gels (Biorad Miniprotean TGX, Biorad, Munich, Germany) and transferred to nitrocellulose membranes. The membranes were blocked with 5% BSA in TBS-Tween and incubated overnight at 4°C with primary antibodies, 1:1500 for anti-Phospho tau S262 (Thermofisher, Waltham, MA, USA), anti-Phospho tau S199 S202 (Thermofisher, Waltham, MA, USA) and anti-Phospho tau T212 S214 (Thermofisher, Waltham, MA, USA), 1:2000 for anti-Phospho tau S202 T205 (Thermo Scientific, Waltham, MA, USA), and 1:6000 anti-total tau (DAKO Agilent, Santa Clara, CA, USA). After three washes with TBS-Tween, membranes were incubated for two hours with Li-Cor fluorescent antibodies at 1:10000 and then washed three additional times with TBS-Tween, once with TBS and then developed in a Li-Cor Odyssey CLx system (Biotech, Bad Homburg, Germany).

### Enzyme linked immunosorbent assay

96-well high binding plate polystyrene microtiter plates (Corning, Tewksbury, MA, USA) were coated with 50 ng/well of aSyn 1 antibody for aSyn detection or 100 ng/well of anti-tau HT7 (MN1000, Thermo Fischer, Rockford, IL, USA) in DPBS and incubated at 4°C overnight. Solution was removed from each well and the wells were washed three times with TBS-Tween. Cell media was added to the wells and left for incubation under shaking at room temperature for either two hours (for aSyn) or overnight after 4 hours of blocking with 3% BSA (for tau).

After the incubation, three washes with TBS-Tween were performed and either anti-αβγ-synuclein (Santa Cruz Biotechnology, Dallas, TX, USA) at a 1:200 dilution or anti-tau (DAKO Agilent, Santa Clara, CA, USA) at a 1:11000 dilution were added to the wells and incubated with shaking for one hour. Next, the wells were washed five times and an anti-rabbit HRP conjugated detection antibody (GE Healthcare, Helsinki, Finland) was added at a 1:5000 dilution. After incubation for one hour, the wells were washed five times and the K-blue aqueous substrate (TMB) was used as a substrate for HRP. After adding the TMB substrate, the reaction was stopped using 1 M H_2_SO_4_. The plates were measured using a Tecan Infinite M200 plate reader at 450 nm.

### Immunofluorescence analyses

Cultures used for immunofluorescence were grown on glass coverslips and fixed with 4% paraformaldehyde (PFA) for 20 minutes. Cells were subsequently permeabilized with 0,1% Triton X-100 at room temperature for 20 minutes. The cells were then blocked with 1,5% bovine serum albumin (BSA, NZYTech, Lisbon, Portugal) for two hours. After blocking cells were incubated overnight at 4°C with the corresponding primary antibodies diluted in 1,5% BSA. The antibodies used were anti-D1,D2-dynamin (1:2000, BD Biosciences, San Jose, CA, USA), anti-D1 dynamin (1:2000, Abcam, Cambridge, UK), aSyn 1 (1:2000, BD Biosciences, San Jose, CA, USA) and anti-tau (1:4000, DAKO Agilent, Santa Clara, CA, USA). The cultures were then washed three times with PBS and then incubated for two hours with the following fluorescent secondary antibodies: Alexa Fluor 568-conjugated donkey anti mouse (1:2000, Life Technologies-Invitrogen, Carlsbad, CA, USA), Alexa Fluor 488-conjugated donkey anti rabbit (1:2000, Life Technologies-Invitrogen, Carlsbad, CA, USA), Alexa Fluor 568-conjugated donkey anti rabbit (1:2000, Life Technologies-Invitrogen, Carlsbad, CA, USA)and Alexa Fluor 488-conjugated donkey anti mouse (1:2000, Life Technologies-Invitrogen, Carlsbad, CA, USA). Finally, cells were stained with DAPI (Roth, Karlsruhe, Germany) for 5 minutes, and coverslipped with Mowiol (Calbiochem, Darmstadt, Germany).

### RT-QuiC assay

Real-time quaking induced conversion (RT-QuiC) was performed as previously reported ^70^. In brief, 100 μl of reaction mixtures were pipetted in black 96-well plates per triplicate (Corning Incorporated, Washington, USA). Composition of the reaction mixtures was: 150 mM NaCl, 1 mM EDTA, 10 μM ThioT, 70 μM SDS, and either 10 μg/ml of monomeric aSyn or 7 µg/ml of monomeric K18 in PBS buffer (pH = 7.1). Just before starting amplification, either anle138b, EGCG or their corresponding vehicles (DMSO and water) were added to the mixtures containing aSyn to a final concentration of 0.1 µM for anle138b and 10 nM for EGCG; for the wells containing K18, fulvic acid was added to a final concentration of 37µM or vehicle (methanol) was added in the same volume.

### Homogeneous time-resolved fluorescence protein aggregation assay

HTRF experiments were performed with the Cis-bio alpha-synuclein aggregation kit and the Cis-bio tau aggregation kit (Cis-bio GmbH, Berlin, Germany) following the manufacturer’s indications. In brief, 5 µl of each conjugated antibody dilution were added to 10 µl of cell lysates with a total protein concentration of 25 ng/ml. The mixture was incubated for 20 hours at 20°C and read in a Tecan Spark 20M plate reader at 665 nm and 620 nm to obtain the ratio of the time resolved FRET (665 nm) and the donor emission (620 nm) that is artifact free.

### Statistical analyses

All data were obtained from at least three independent experiments and were expressed as mean values ± standard deviation (SD), determined as 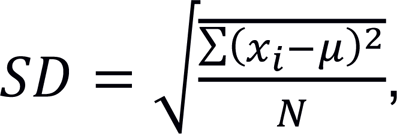 where *N* is the number of experiments performed, *x_i_* each value obtained and *μ* the mean value. Two-group comparisons were performed with Student’s t test. Differences were considered as statistically significant at p < 0,05. Statistical analyses were performed in Excel (Microsoft, Seattle, WA, USA).

## Acknowledgements

This work has received support from the EU/EFPIA/Innovative Medicines Initiative 2 Joint Undertaking (IMPRIND grant n° 116060). The opinions expressed and arguments employed herein do not necessarily reflect the official views of these funding bodies. ADM is supported by a postdoctoral fellowship from the Galician Government (Programa de axuda á etapa posdoutoral, XUGA, GAIN, ED481B 2017/053). M.Z. was supported by the advanced grant ‘787679 - LLPS-NMR’ of the European Research Council.

## Author contributions

TFO, ADM and EV designed the study. ADM performed cell cultures, treatments, prepared samples, flow cytometry, immunoblotting, ELISA, immunofluorescence, RT-QuiC, HTRF and statistical analyses. AK and MSCO prepared the recombinant proteins. TFO, AL, SR, CG and MZ contributed material/reagents/analysis tools. ADM and EV prepared all figures. ADM and TFO wrote the manuscript. All authors discussed the results and contributed to the manuscript.

## Competing interests

CG is co-inventor in a patent application related to the anle138b compound included in this study. CG is shareholder and co-founder of MODAG GmbH. AL is partly employed by MODAG. All other authors declare no competing interests.

---

**Supplementary Fig. S1.**
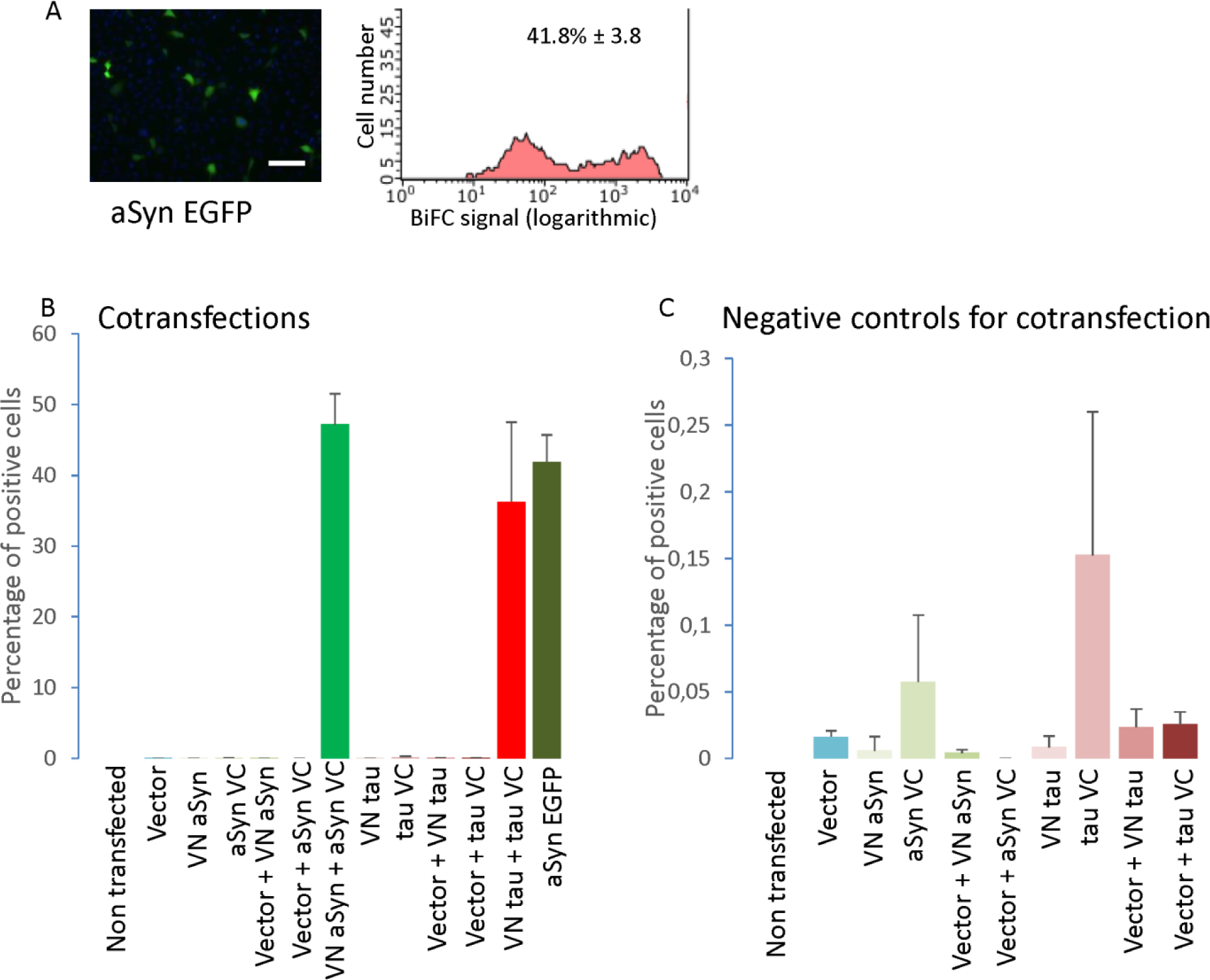
Both halves of venus are needed for the reconstitution of the fluorophore. A. Representative pictures and histogram of the flow cytometry for aSyn EGFP showing similar results to the ones obtained in cotransfection. Cells with a value of fluorescence intensity above 120 random fluorescence units are considered positive. Scale bar 200 µm. B. Co-transfection combinations show percentages of positive cells above 30%C. Negative controls show less than 0.2% positive cells. Error bars represent SD.

**Supplementary Fig. S2.**
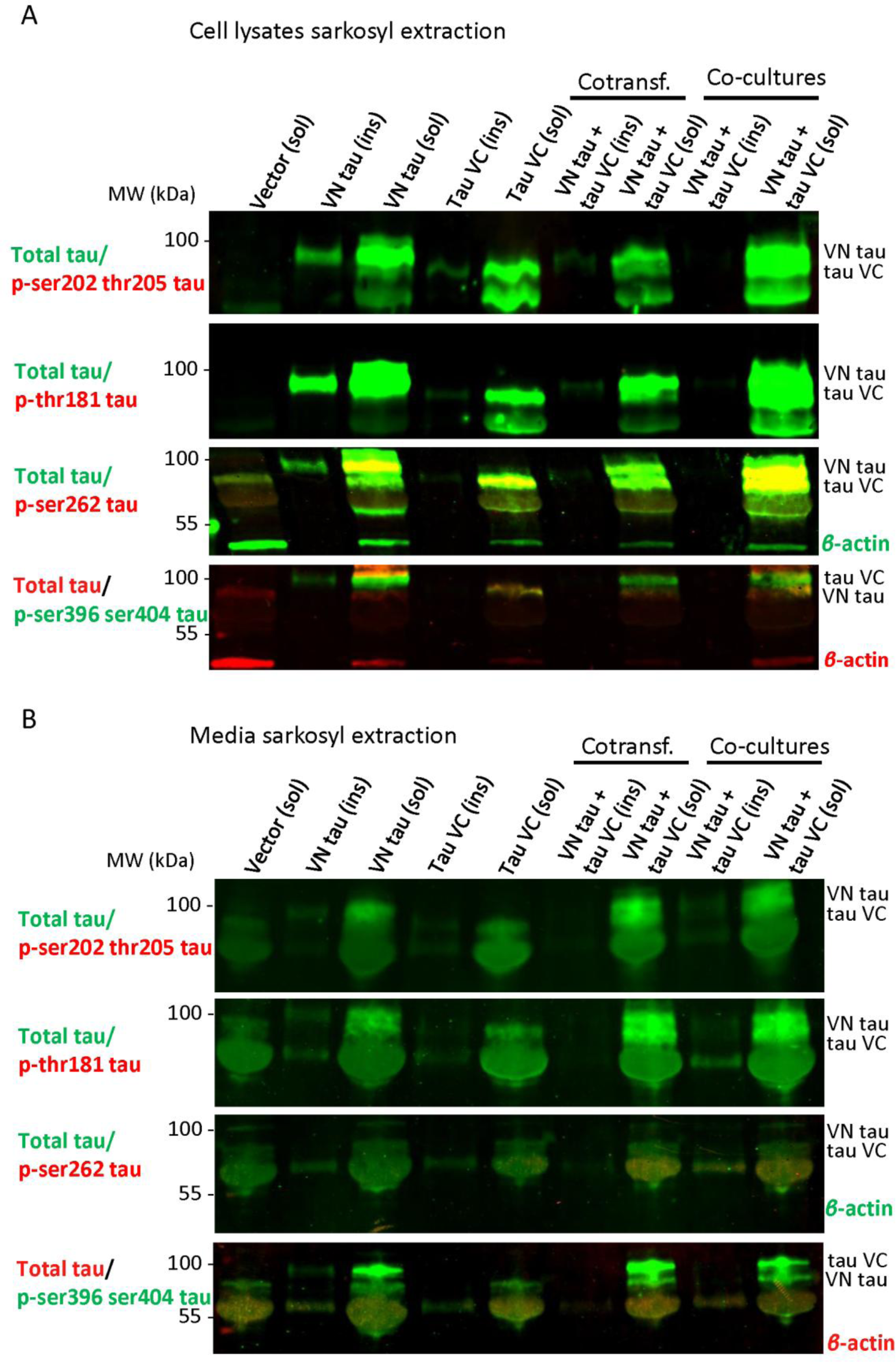
Tau is secreted in sarkosyl soluble and insoluble fractions. A. Western blot showing different tau BiFC fragments in cotransfected cells and co-cultured cells in sarkosyl soluble (sol) and insoluble fractions (ins) and phosphorylation of serine 262, serine 396 and serine 404. Full length blots are presented in Supplementary Figure 6F-I. B. Western blot showing different tau BiFC fragments, alone, cotransfected and in co-culture, present in cell media in the sarkosyl soluble and insoluble fractions. Full length blots are presented in Supplementary Figure 6J-M.

**Supplementary Fig. S3.**
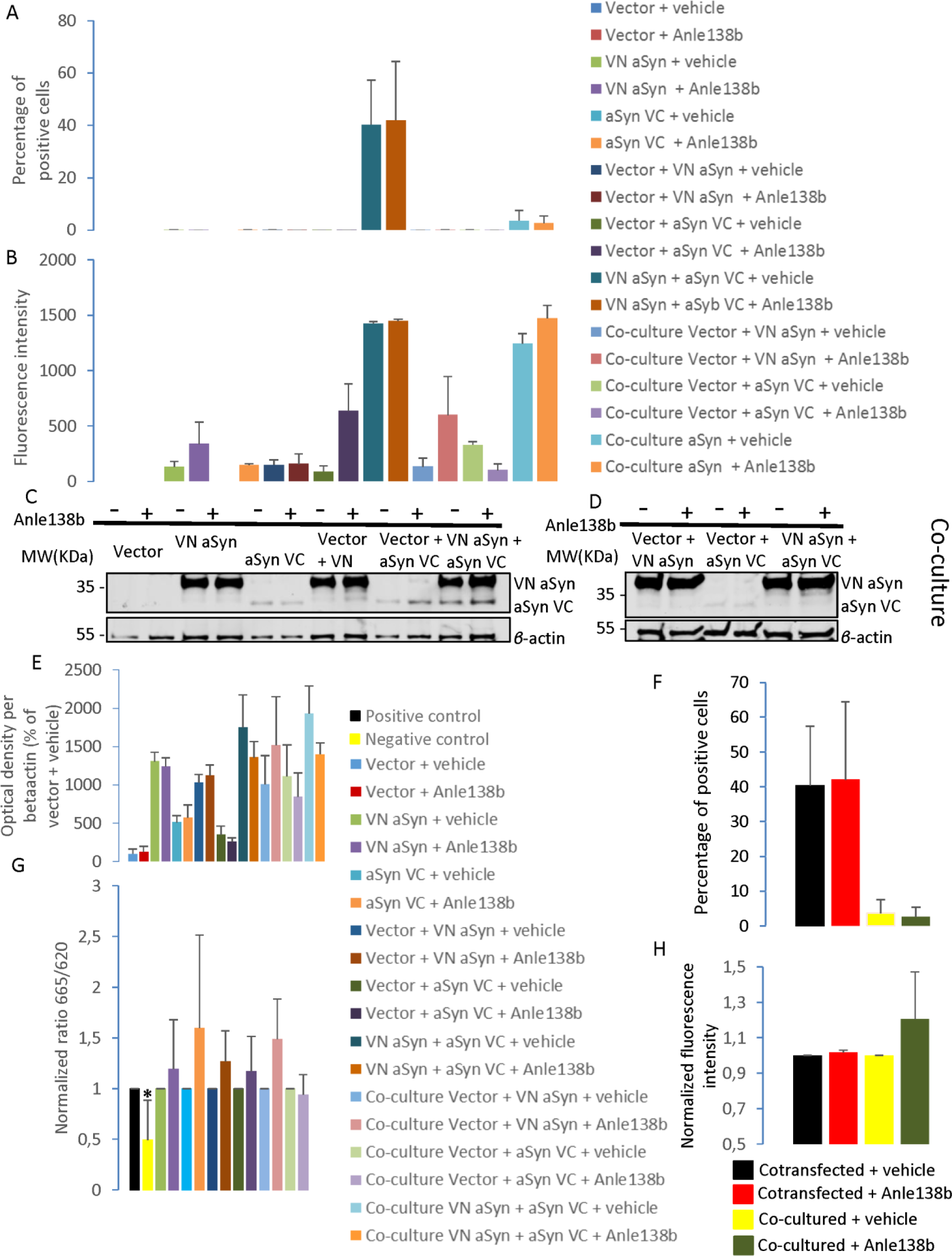
Treatment with anle138b inhibits aSyn aggregation without altering protein expression. A. Flow cytometry results show that treatment with anle138b does not lead to a significant decrease in the percentage of positive cells. B. Flow cytometry results showing fluorescence intensity levels obtained in cells treated with vehicle and anle138b. C. Western blots of transfected and non-co-cultured cells treated and non-treated with anle138b. Full length blots are presented in Supplementary Figure 6N. D. Western blots of transfected and co-cultured cells treated with anle138b. Full length blots are presented in Supplementary Figure 6O. E. Western blot results of normalized optical density per beta-actin obtained in cells treated with vehicle and Anle138b showing that treatment with Anle138b does not lead to significant changes in protein expression (N=4). F. Flow cytometry results showing the percentage of positive cells in co-transfected cells and in co-cultured cells. G. HTRF results for the normalized ratio 665/620 in cells treated with vehicle and anle138b. Treatment with anle138b does not lead to a significant decrease in protein aggregation levels 12 hours after administration in comparison with cells treated with vehicle. Positive control shows a significantly higher value than the negative control. (N=4) H. Flow cytometry results for the normalized fluorescence intensity show no statistically significant differences (N=3). * p< 0.05 in comparison with the control. $ p<0.05 in comparison with monomeric aSyn + vehicle. Error bars represent SD.

**Supplementary Fig. S4.**
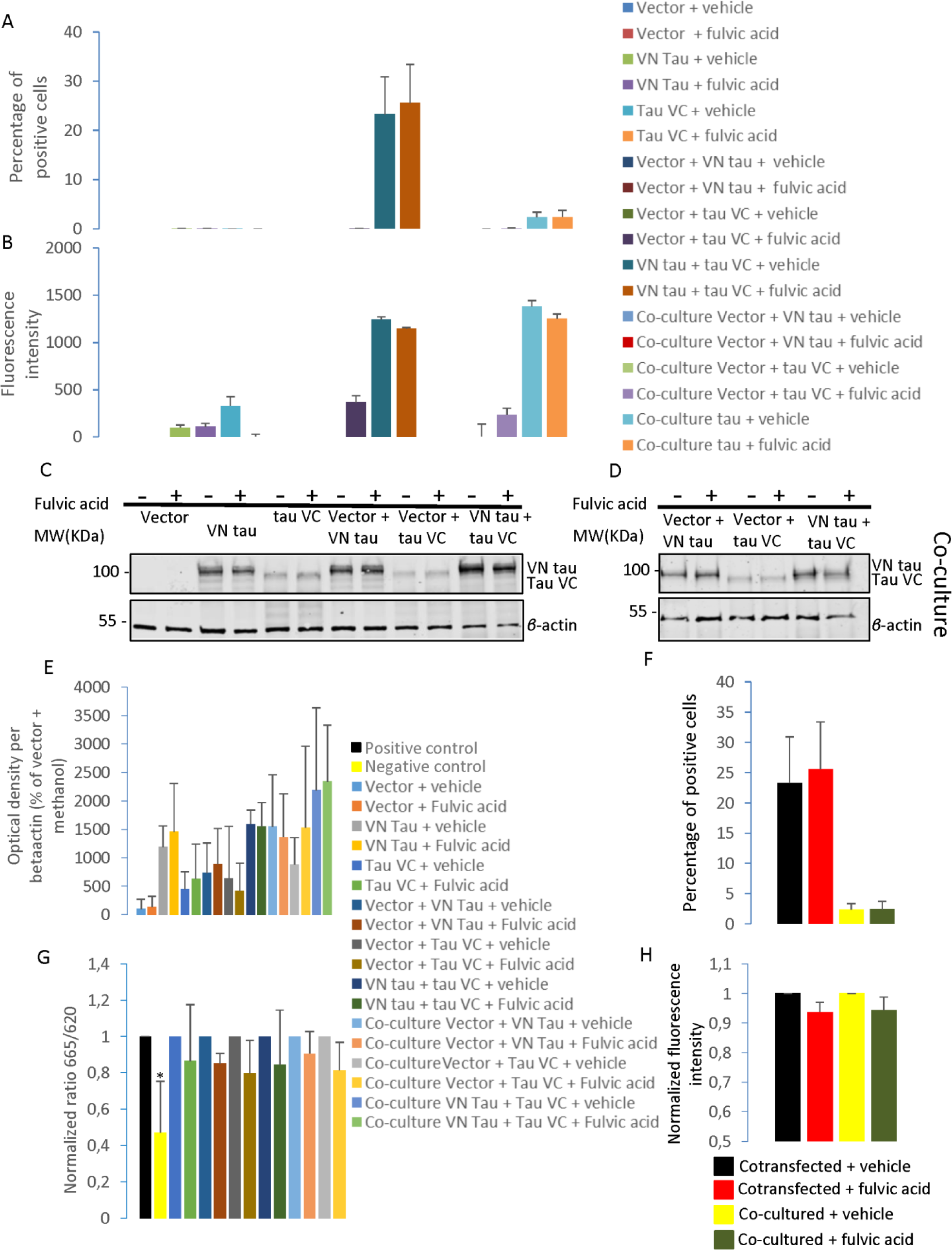
Treatment with fulvic acid inhibits tau aggregation without altering protein expression. A. Flow cytometry results showing the percentage of positive cells when treated with vehicle and fulvic acid. Treatment with fulvic acid does not lead to a significant decrease in the percentage of positive cells. B. Flow cytometry levels of fluorescence intensity levels obtained in cells treated with vehicle and fulvic acid. C. Western blot of transfected and non-co-cultured cells treated and non-treated with fulvic acid. Full length blots are presented in Supplementary Figure 6P. D. Western blot of transfected and co-cultured cells treated with fulvic acid. Full length blots are presented in Supplementary Figure 6Q. E. Western blot quantification of the normalized optical density per beta-actin obtained in cells treated with vehicle and fulvic acid showing no significant differences between cells treated with vehicle or with fulvic acid (N=3). F. Flow cytometry results showing the percentage of positive cells in co-transfected cells and in co-cultured cells. G. HTRF values obtained for the normalized ratio 665/620 in cells treated with vehicle and fulvic acid. Positive control shows a significantly higher value than negative control (N=4). H. Flow cytometry results for the normalized fluorescence intensity show no statistically significant differences (N=3)* p< 0.05 in comparison with the control. $ p<0.05 in comparison with monomeric tau K18 + vehicle. Error bars represent SD.

**Supplementary Fig. S5.**
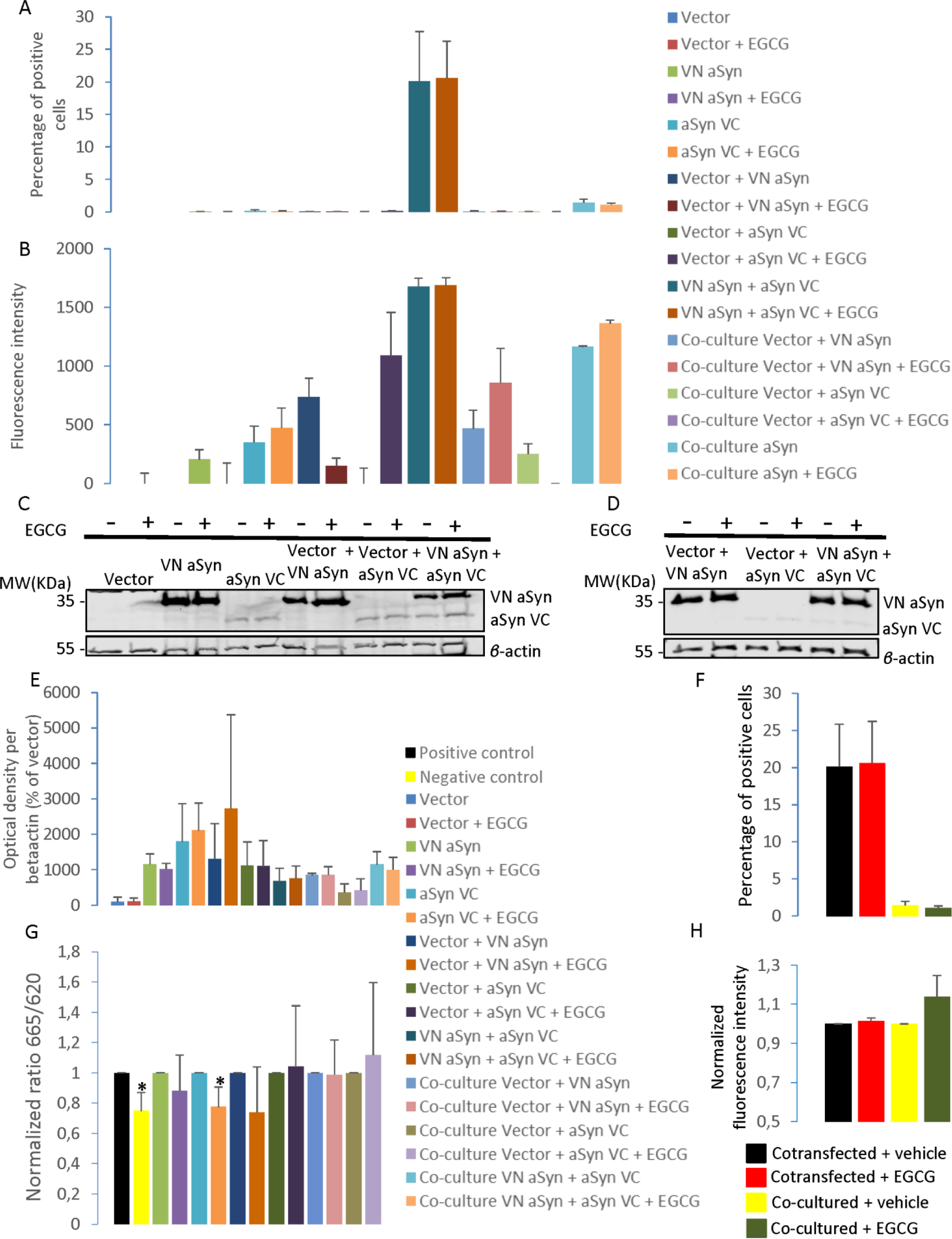
Treatment with EGCG inhibits aSyn aggregation without altering protein expression. A. Flow cytometry results showing the percentage of positive cells treated with EGCG. Treatment with EGCG does not lead to a significant decrease in the percentage of positive cells. B. Flow cytometry levels of fluorescence intensity obtained in cells treated and untreated with EGCG. Treatment with EGCG does not lead to a significant decrease in fluorescence intensity. C. Western blot of transfected and non-co-cultured cells treated and non-treated with EGCG. Full length blots are presented in Supplementary Figure 6R. D. Western blot of transfected and co-cultured cells treated with EGCG. Full length blots are presented in Supplementary Figure 6S. E. Western blot quantification of the normalized optical density per beta-actin obtained in cells treated and untreated with EGCG showing no significant differences between treated and untreated cells (N=3). F. Flow cytometry results showing the percentage of positive cells in co-transfected cells and in co-cultured cells. G. HTRF measurements of the normalized ratio 665/620 in cells treated and untreated with EGCG. The positive control shows a significantly higher value than the negative control (N=4). H. Flow cytometry results for the normalized fluorescence intensity show no statistically significant differences (N=3).* p< 0.05 in comparison with the control. Error bars represent SD.

**Supplementary Fig. S6.**
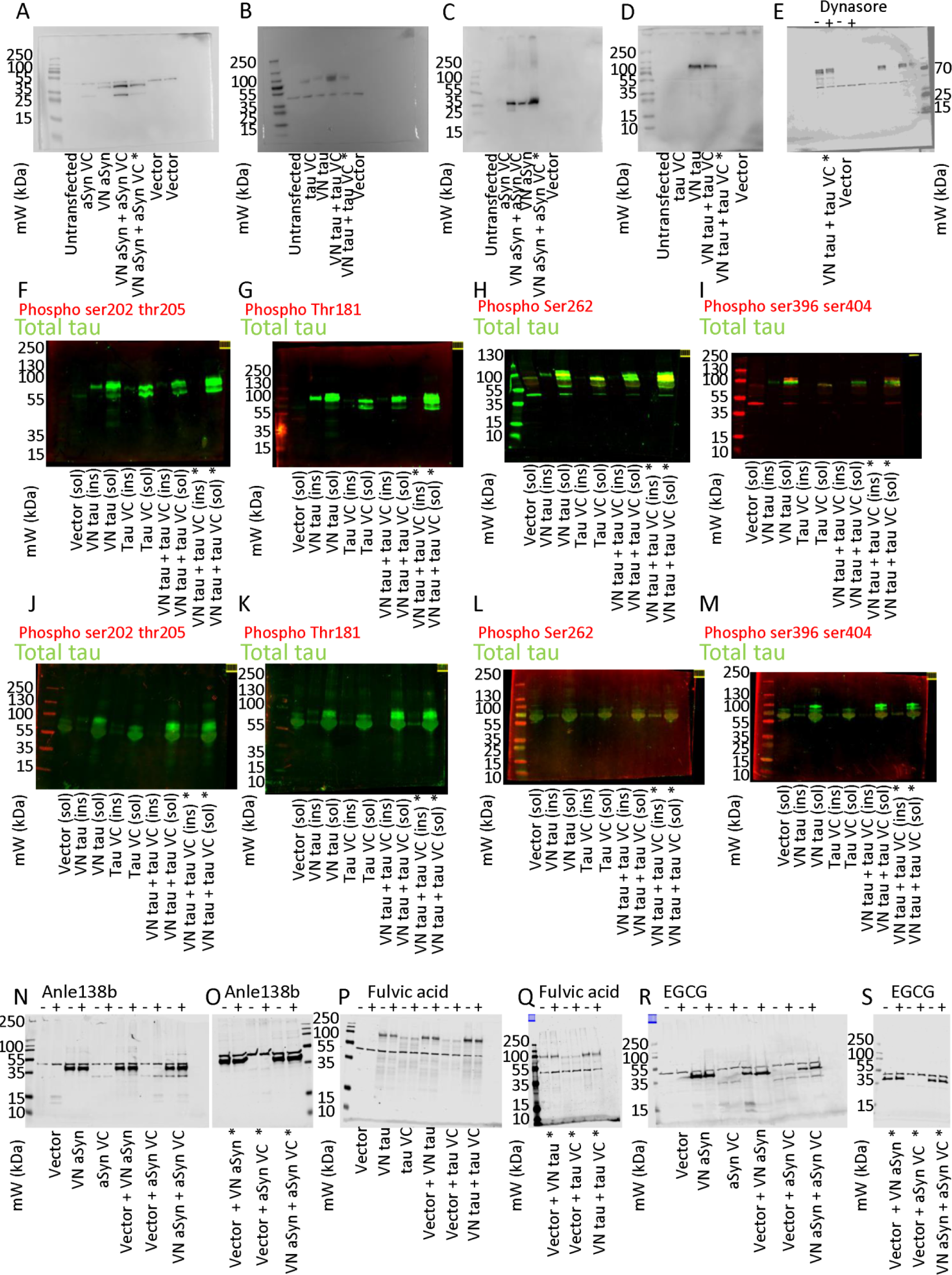
Full length blots. A. Full lengh blots of the cropped image from Fig. 1E. B. Full length blots of the cropped image from Fig. 1F. C. Full length blots of the cropped image from Fig. 2A. D. Full length blots of the cropped image from Fig. 2B. E. Full length blots of the cropped image from Fig. 4H. Image is presented as an overlay of the full blot blot and the marker. F-I. Full length blots of the cropped images from Supplementary Figure 2A. J-M. Full length blots of the cropped images from Supplementary Figure 2B. N. Full length blots of the cropped images from Supplementary Fig.S3C. O. Full length blots of the cropped images from Supplementary Figure 3D. P. Full length blots of the cropped images from Supplementary Figure 4C. Q. Full length blots of the cropped images from Supplementary Figure 4D. R. Full length blots of the cropped images from Supplementary Figure 5C. S. Full length blots of the cropped images from Supplementary Figure 5D. Asterisks (*) indicate co-cultured cells.

## Notes

#### Summary of Updates

Authors updated. Minor changes in the introduction. Supplemental files updated.

